# A robust method of extraction and GC-MS analysis of Monophenols exhibited UV-B mediated accumulation in Arabidopsis

**DOI:** 10.1101/2021.07.30.454506

**Authors:** Maneesh Lingwan, Shyam Kumar Masakapalli

**Affiliations:** BioX Center, School of Basic Sciences, Indian Institute of Technology Mandi, Kamand - 175075, Himachal Pradesh, India

**Keywords:** Plant Phenolics, MTBE extraction, GC-MS, metabolite profiling, Monophenols, UV-B, Arabidopsis

## Abstract

Studies on specialised metabolites like phenolics are of immense interest owing to their significance to agriculture, nutrition and health. In plants, phenolics accumulate and exhibits spatial and temporal regulations in response to growth conditions. Robust methodologies aimed at efficient extraction of plant phenolics, their qualitative and quantitative analysis is desired. We optimised the analytical and experimental bottlenecks that captured free, ester, glycoside and wall-bound phenolics after acid or alkali treatments of the tissue extracts and subsequent GC-MS analysis. Higher recovery of phenolics from the methanolic extracts was achieved by through a) Ultrasonication assisted extraction along with Methyl tert-butyl ether (MTBE) enrichment b) nitrogen gas drying and c) their derivatisation using MSTFA for GC-MS analysis. The optimised protocol was tested on Arabidopsis rosette exposed to UV-B radiation (280-315 nm) which triggered enhanced levels of 11 monophenols and might be attributed to photoprotection and other physiological roles. Interestingly, coumaric acid (308 m/z) and caffeic acid (396 m/z) levels were enhanced by 12-14 folds under UV-B. Other phenolics such as cinnamic acid (220 m/z), hydroxybenzoic acid (282 m/z), vanillic acid (312 m/z, gallic acid (458 m/z), ferulic acid (338 m/z), benzoic acid (194 m/z), hydroxycinnamic acid (368 m/z) and protocatechuic acid (370 m/z) also showed elevated levels by about 1 to 4 folds. Notably, vanillin (253 m/z) was detected only in the UV-B exposed tissues. The protocol also comprehensively captured the variations in the levels of ester, glycoside and wall-bounded phenolics with high reproducibility and sensitivity. The robust method of extraction and GC-MS analysis can readily be adopted for studying phenolics in plant systems.

## 1. Introduction

In plants, metabolism can be divided into primary or secondary depending on the pathways they support for growth, development or other specialised tasks (Aharoni and Galili, 2011)(Fang et al., 2019). While primary metabolism is universal, secondary metabolism is often unique to specific plant families. Metabolites are small molecules that facilitate metabolic processes. Primary metabolites like amino acids, sugars, organic acids and others involved in processes such as photosynthesis, respiration etc. (Fang et al., 2019) Metabolites like alkaloids, terpenoids, phenolics, anthocyanins, coumarins and others support secondary metabolism leading to an array of phytochemicals that play specific roles like protection under stress. Among these the phenolics, confer chemical-biological roles supporting growth, development and protection under stress (Bueno et al., 2012)(Singh et al., 2021). Accumulation of phenolics is spatial, temporal and stimuli regulated and can be specific to tissue types, species, and developmental stages (Barros et al., 2012)(Lingwan et al., 2020). The analysis of phenolics is essential to understand the adaptation mechanisms in plants. The plant phenolics can broadly be classified into polyphenols (lignin, tannins), oligophenols (coumarins, stilbenes, flavonoids) and monophenols. The chemical structure of phenolic compounds has a phenolic ring with one or more hydroxyl groups, which are reported to promote plant defence, enhance antioxidants by scavenging free radicals. The levels of phenolics accumulation also gets modulated due to targeted or non-targeted engineering of plants (Escarpa and Gonzalez, 2001)(Parr and Bolwell, 2000).

Monophenols are categorized into two important classes cinnamic acid or benzoic acid derivatives which serve as precursors for caffeic acid, coumaric acid, ferulic acid, protocatechuic acid, hydroxybenzoic acid, vanillic acid, gallic acid and others (Marchiosi et al., 2020). These monophenols accumulate in various plant tissues as either free or wall-bound in the form of esters or glycosides and are reported to play a significant role in physiology and anatomy **(Fig. 1)**. Several of these monophenols owing to their high antioxidant and radical scavenging abilities has led to identifying their applications to the food and health industry (Shahidi and Yeo, 2018)(Anantharaju et al., 2016). Phenolics are reported to provide health benefits by conferring protection against cardiovascular, cancer and inflammatory diseases. Considering all the advantages, qualitative and quantitative profiling of the phenolics in plant tissues (edible and non-edible) are crucial so that their roles can appropriately be investigated (Shahidi and Ambigaipalan, 2015)(Balasundram et al., 2006).

**Fig. 1.**
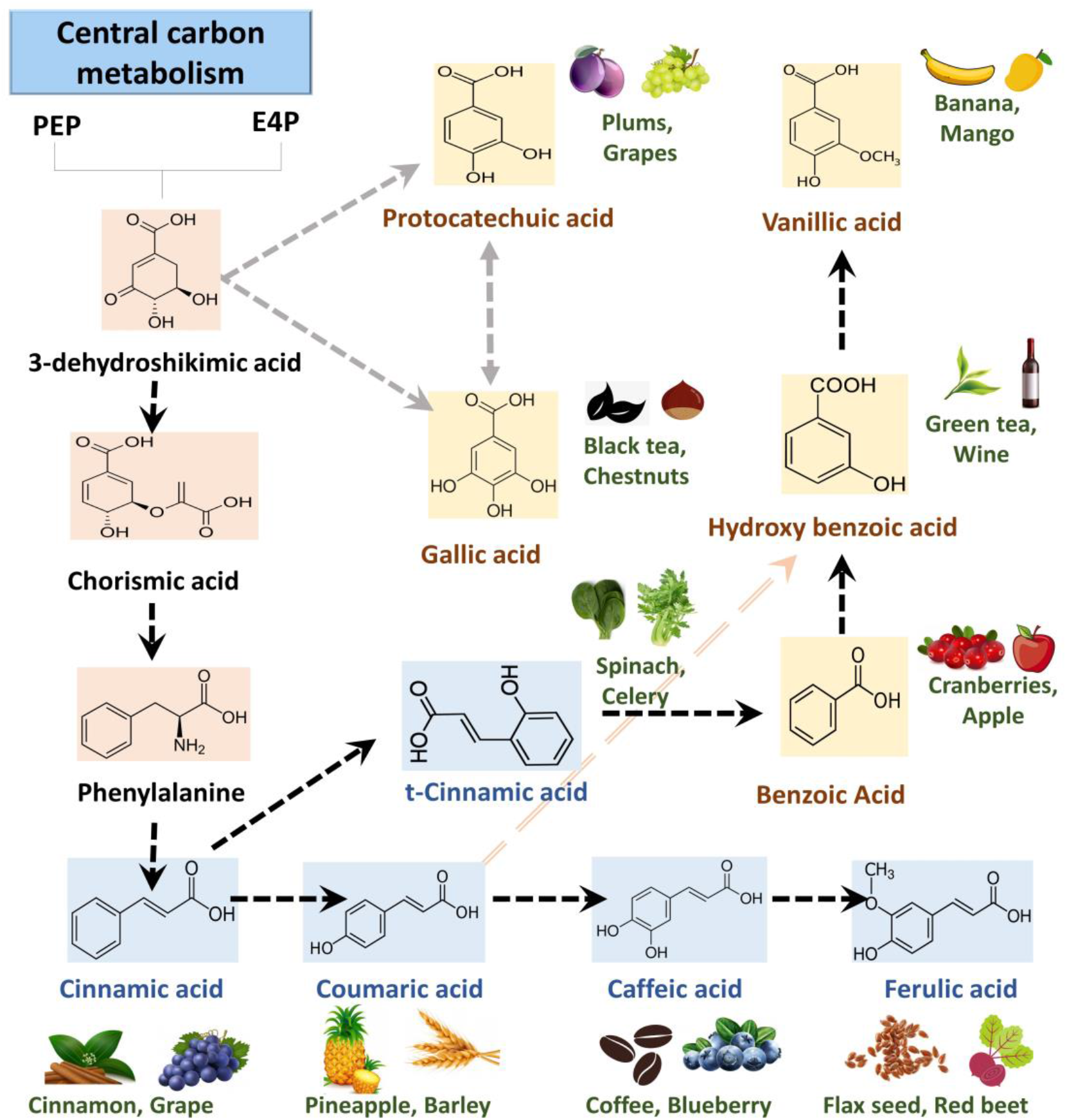
Metabolic pathway and chemical structure of plant phenolics. Monophenols such as benzoic acid, cinnamic acid and their derivatives are biosynthesized via the shikimate pathway. These phenolics remains in free form or accumulated as ester bound and glycoside bound performing various physiological roles. Phenolics are known to be present in fruits, vegetables etc. thereby playing role in health and nutrition. Benzoic acid derivatives are in yellow and cinnamic acid derivatives are in blue background.

Profiling of the phenolics from plant tissues has been widely studied but there is further needed to develop robust methods with higher sensitivity. The variations in the range of physicochemical properties, molecular masses and structure among classes of phenolics often limits the profiling. Separation and identification of several phenols simultaneously are highly challenging and often need optimisation and development of suitable methods (Lingwan et al., 2020)(Lin et al., 2007)(Lisec et al., 2006). Gas chromatography-mass spectrometry (GC-MS) can be a sensitive, fast and reproducible platform for the determination of phenols as will be demonstrated in this work. For GC-MS analysis of phenolics from plant tissues, we need to first extract them, derivatise and identify them based on their m/z fragments. Methods that enhance the limit of detection (LOD), targeted enrichment, separation, derivatisation and data analysis pipelines will be widely useful for targeted and non-targeted analysis of phenolics. In this work, a protocol involving steps like ultrasonication, Methyl tert-butyl ether (MTBE) solvent extraction and derivatisation has been optimised for subsequent analysis of phenolics using GC-MS qualitatively and quantitatively. The robust protocol allowed the detection of free, ester bound, glycoside bound and wall-bound phenolics with high sensitivity. The application of the protocol in plant research has been demonstrated by profiling phenols in *Arabidopsis thaliana* tissues under normal and perturbed conditions (UV-B radiation) that affect levels of phenolics

## 2. Experimental Procedure

### 2.1. Reagents and chemicals

Moorashige & Skooge (MS) medium, 2-N-morpholino-ethanesulfonic acid (MES), sucrose, phyta agar, sodium hypochlorite, aluminium foil, forceps were purchase from Himedia labs. Methanol, Ribitol, Methyl tert-butyl ether (MTBE), Pyridoxine hydrochloride, Methoxamine hydrochloride (MeOX), N-Methyl-N-trimethylsilyl trifluoroacetamide (MSTFA), O-bis(trimethylsilyl)trifluoroacetamide (BSTFA), Hydrochloric acid (HCl), Sodium dodecyl sulphate (SDS), Sodium thiosulphate, Ethanol, Acetone, Sodium hydroxide (NaOH) and Commercial Phenol standards as Cinnamic acid, p-Hydroxybenzoic acid, Vanillin, Vanillic acid, p-Coumaric acid, Gallic acid, Ferulic acid and cis-Caffeic acid were purchase from Sigma-Aldrich. All the reagents were checked for analytical grade and safely stored at recommended temperature.

### 2.2. Equipment and Software

pH meter, Autoclave, Petri plates, Percival LED22C8 growth cabinet, Lyophilizer, Ice bucket, Thermomixer (Eppendorf ThermoMixer C), Digital ultrasonicator, UV light meter (Sper Scientific), Centrifuge, Speed Vac drier (DNA Speedvac Thermo Fisher Scientific), GC-MS, in software Chemstation, Metalign, NIST 17, Fiehn Metabolomics library, Metaboanalyst 5.0, GraphPad Prism 8 and Microsoft Excel was used.

### 2.3. Preparation of media and growth conditions for Arabidopsis

Columbia (Col-0) ecotype of *A. thaliana* seeds was sterilized in 5% sodium hypochlorite for 4 minutes and washed in autoclaved water at least 5 times. The seeds were inoculated on sterile plates containing MS media (with 0.05% MES, 1% sucrose, 0.8% agar, pH-5.78) and kept at 4 °C for 48 hours in darkness for stratification. The plates were transferred to a plant growth chamber (Percival LED22C8) and incubated at 22°C, 70% humidity with a 16/8-hour light-dark cycle. The intensity of visible light was set to 110 µmol/m2/s and for UV-B treatment, 1.4 W/m2 UV-B light intensity was used and measured by UV light meter. For UV-B treatment, on the 14th day, Arabidopsis rosette were exposed to 16 hours of UV-B lights. After UV-B treatments, rosette was harvested from independent (n=4) plates using liquid nitrogen and crushed with the help of mortar and pestle. The crushed powder is lyophilized overnight to remove all moisture and stored at -20°C until further use (Yadav et al., 2019a)(Yadav et al., 2019b).

### 2.4. Preparation of extraction solvents and standards

The extraction solvents as MTBE, Methanol and Water were kept back at -20°C before extraction. The commercial phenol standards were dissolved in 80% methanol at a stock of 100 µg/ml and stored at -20°C until use. The working dilution range for the standard is started from 10µg /ml. Ribitol in dilution of 10 µg/ml was used as internal standards for GC-MS analysis. Phenolics standards of cinnamic acid, hydroxybenzoic acid, vanillin, vanillic acid, coumaric acid, gallic acid, ferulic acid and cis-caffeic acid were taken for quantification.

### 2.5. Phenolics extraction

Lyophilized tissue powder (∼20 mg) is weighed in a screw cap vial (2ml) and 600 ul of ice-cold methanol was added and vortexed for 2 minutes. The vials were incubated in a shaker at 4 °C at 950 rpm for 30 minutes followed by incubation again in a thermo shaker at 70°C at 950 rpm for 5 minutes. Once cooled on ice, 200 ul methanol, 150 ul water and 50 ul ribitol (10 µg /ml) as an internal standard was added. The last incubation step was repeated and then ultra-sonicated (40±3 kHz frequency) for 5 minutes. After sonication, the vials were mixed at room temperature in a thermoshaker at 950 rpm for 5 minutes. The extracts were then centrifuged at 15000 g for 15 min at room temperature. 30 µl supernatant was collected in a new 1.5 ml eppendorf tube and dried under-speed vacuum drier and subjected to derivatization for GCMS analysis **(Fig. 2)**. The aliquots (250 ul each) from the remaining supernatant were used for extraction of free soluble, glycoside bound and ester bound phenolic as detailed below. The pellets were further used to extract wall-bound phenolics as detailed below. All samples were stored at -20°C till further use.

**Fig. 2.**
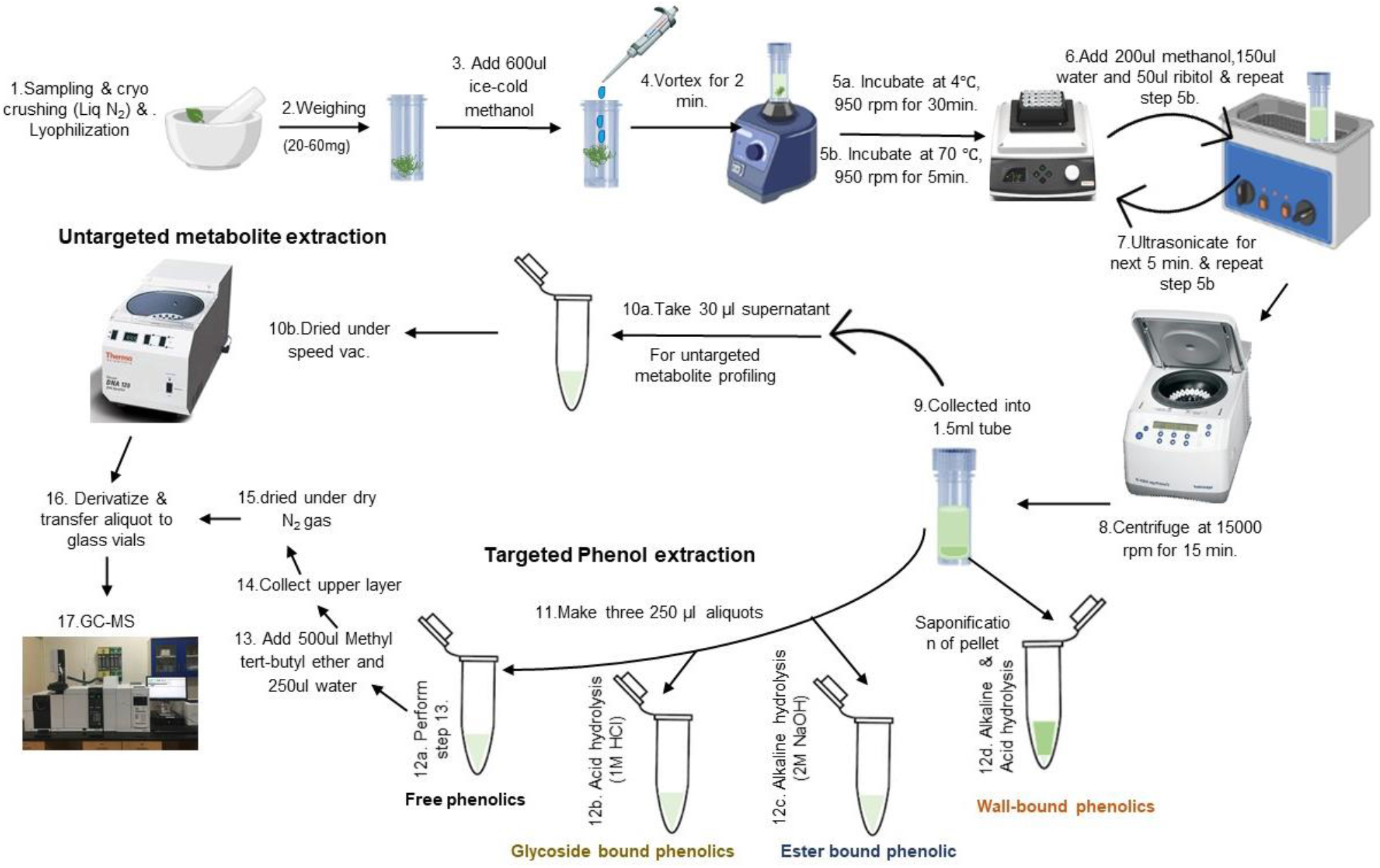
A schematic workflow for robust extraction and analysis of free and bound phenolics in Arabidopsis. The major steps involve extraction (steps 1-8), fractionation and treatment for targeted phenol extraction (steps 9-14), drying and GC-MS analysis (steps 15-17).

#### 2.5.1. Extraction of free soluble phenolics

For free phenolic extraction, an MTBE method was optimised and used. To 250 µl of supernatant aliquot, 500 µl of chilled cold (−20 oC) MTBE and 250 µl of milli-Q water is added and allowed to stand for few minutes until clear phase separation appears. The upper layer (∼500 µl) was collected in fresh Eppendorf tube and was completely dried under N_2_ gas. The dried samples were subjected to derivatisation and GC-MS analysis.

#### 2.5.2. Extraction of glycoside-bound phenolics

The glycoside-bound phenolics were extracted as suggested by (Ascensao et al., 2003) with slight modifications. To 250 µl of supernatant aliquot, 25 µl of 1M HCl were added to the vial and left for acid hydrolyses at 96 oC for 60 minutes. After acid hydrolysis, chilled 500 µl of MTBE and 250 µl of milli-Q water were added. Then upper layer was collected, completely dried under dry N_2_ gas, derivatised and analysed in GC-MS.

#### 2.5.3. Extraction of ester-bound phenolics

The ester-bound phenolics were extracted as suggested by (Cvikrová et al., 1993) with slight modifications. To 250 µl of supernatant aliquot, 62.5 µl of 1M NaOH was added and incubated for 180 minutes at room temperature for alkaline hydrolysis. Then 20 µl of 1M HCl was added and mixed properly. After mixing, 500 µl of MTBE and 250 µl of milli-Q were added and the upper layer was collected. Then collected layer was dried under dry N_2_ gas, further derivatised and analysed in GC-MS.

#### 2.5.4. Extraction procedure for wall-bound phenolics

The wall-bound phenolics were extracted as suggested by (Campbell and Ellis, 1992) with slight modifications. The tissue pellet was dried at 70°C for 24 hours. Then saponification was performed by treating the pellet with 1.5% (w/v) SDS and 5 mM sodium thiosulphate for 20 min at 26°C. Sample was centrifuged at 5000g for 15 min. Then pallet was washed safely with boiling ethanol to remove any alcohol soluble phenolics. Next, pellet was carefully treated with acetone and dried with N_2_ gas. Completing the washing and drying, the dried weight of cell wall material was taken for calculating the amount of wall-bound phenolics. After Completing pre-processing steps, the pellet was subjected to alkaline hydrolysis with 250 µl of 1M NaOH for 24 hours in dark and then acidified with 250 µl of 1M HCl. In the final purification step, 500 ul of Methyl tert-butyl ether and 250 µl of Milli-Q water were added and an upper separated layer was collected, dried under N_2_ gas and further derivatised for GC-MS analysis **(Fig. 2)**.

### 2.7. Gas Chromatography -Mass Spectroscopy for profiling of plant phenols

#### 2.7.1. Sample derivatization for GC-MS

Dried extracts for untargeted profiling and phenolics were subjected to similar derivatisation reactions. All the samples and standards were MeOX-TMS derivatized using pyridine, methoxamine hydrochloride and N-Methyl-N-trimethylsilyl trifluoroacetamide. MeOX-TMS derivatization includes the addition of 35 μl of pyridine containing methoxylamine hydrochloride (20 mg/ml) to the dried sample. The sample was incubated at 37°C, 900 rpm for 2 hours, then 49 μl of MSTFA (N-methyl-N- (trimethylsilyl)-trifluoroacetamide) was added to the sample and re incubated for 30 min. After incubation, sample was centrifuged at 13,000 g for 10 min and the supernatant was transferred to new inserts for GC-MS data acquisition(Shree et al., 2019)(Masakapalli et al., 2014). To examine better derivatization methodology, Samples and standards were also derivatized with O-bis(trimethylsilyl)trifluoroacetamide (BSTFA) to compare the silylation potential of BSTFA with MSTFA for phenols.

#### 2.7.2. GC-MS data acquisition parameters

The data acquisition was performed on GC-MS (GC ALS-MS 5977B, Agilent Technologies) equipped with HP-5ms (5% phenyl methyl siloxane) column (30 m x 250 μm x .25 μm). 1 µl of the derivatized sample was injected in splitless mode. Total run time was set for 50 min with helium used as a carrier gas at a constant flow rate of 0.6 ml/min. The parameter for scan mass range was set from 50 to 600 m/z with an electron ionization of 70 eV. The initial oven programme was set to 50 °C, then it raised to 70 °C in 5 min hold time. The ramp rate was set for 10 °C/min and 5 °C/min with a hold time of 10 min for each to achieved 200 °C and 300 °C temperature respectively (**Table S1)**. All the acquisition parameters were controlled by the chemstation software(Shree et al., 2019)(Saini et al., 2020).

#### 2.7.3. Pre- and post-processing steps in data analysis

The acquired raw file for each condition (n=4) was checked for reproducibility and subjected to baseline correction using Metalign software (Lommen and Kools, 2012). Mass Hunter and Chemstation software were used for profiling and data analysis. For samples and phenols standards, peaks spectra were identified based on the mass ion fragments (m/z), retention time (RT), identifier ions and match against the NIST 17 and Fiehn Metabolomics library. Metabolites having more than 70% probability score in NIST and Fiehn metabolomics library were taken for further analysis (Kopka et al., 2005). The relative peak abundances of phenolics were extracted and subjected to fold changes analysis after normalisation with internal standard. The final metadata excel file was prepared for statistical analysis.

### 2.8. Statistical and multivariate data analysis

A metadata file containing metabolite name and comparative peak abundance were subjected to PCA analysis using Metaboanalyst 5.0. The area of the internal standard ribitol is used to obtain the relative proportions of the peaks which in turn lead to the calculation of fold changes of metabolite/peak levels among the treatments. Basic statistical analysis was used for evaluating the significance or non-significance of data using GraphPad Prism8 (Xia and Wishart, 2016).

## 3. Results

### 3.1. MTBE and ultrasonication assisted extraction showed enriched recovery of phenolics

Ultrasonic assisted extraction (UAE) of plant tissue was reported for improving extraction efficiencies of metabolites (Pan et al., 2012). In our samples extracted under 80% methanol, we observed an additional step of ultrasonication at 40±3 kHz frequency for 5 minutes improved the extraction of phenols and other metabolites. The GC-MS analysis of ultrasonicated samples showed higher peak intensity and better separation **(Fig. 3A)**. Further, MTBE based phase separation of the methanolic extracts showed enriched recovery of phenolics **(Fig. 3B)**. In methanolic extracts, we detected 57 metabolites primarily belonging to sugars, organic acid, amino acid and some phenolics. In MTBE extracts we detected 37 metabolites with higher coverage of phenolics and other metabolites **(Fig. 3C)**. Principal component analysis (PCA) plots revealed that MTBE and methanolic solvent have varied extraction potential for classes of metabolites. PCA plots with scores of 96.9% and 1.4% across the two principal components PC1 and PC2 respectively highlighted the significant differences among the extraction profiles between solvents **(Fig. S1)**. Variable Importance in Projection (VIP) score pattern analysis further highlighted that MTBE has higher efficiency for phenolics, whereas 80% methanol showed higher extraction for sugars and organic acids **(Fig. 3D)**.

**Fig. 3.**
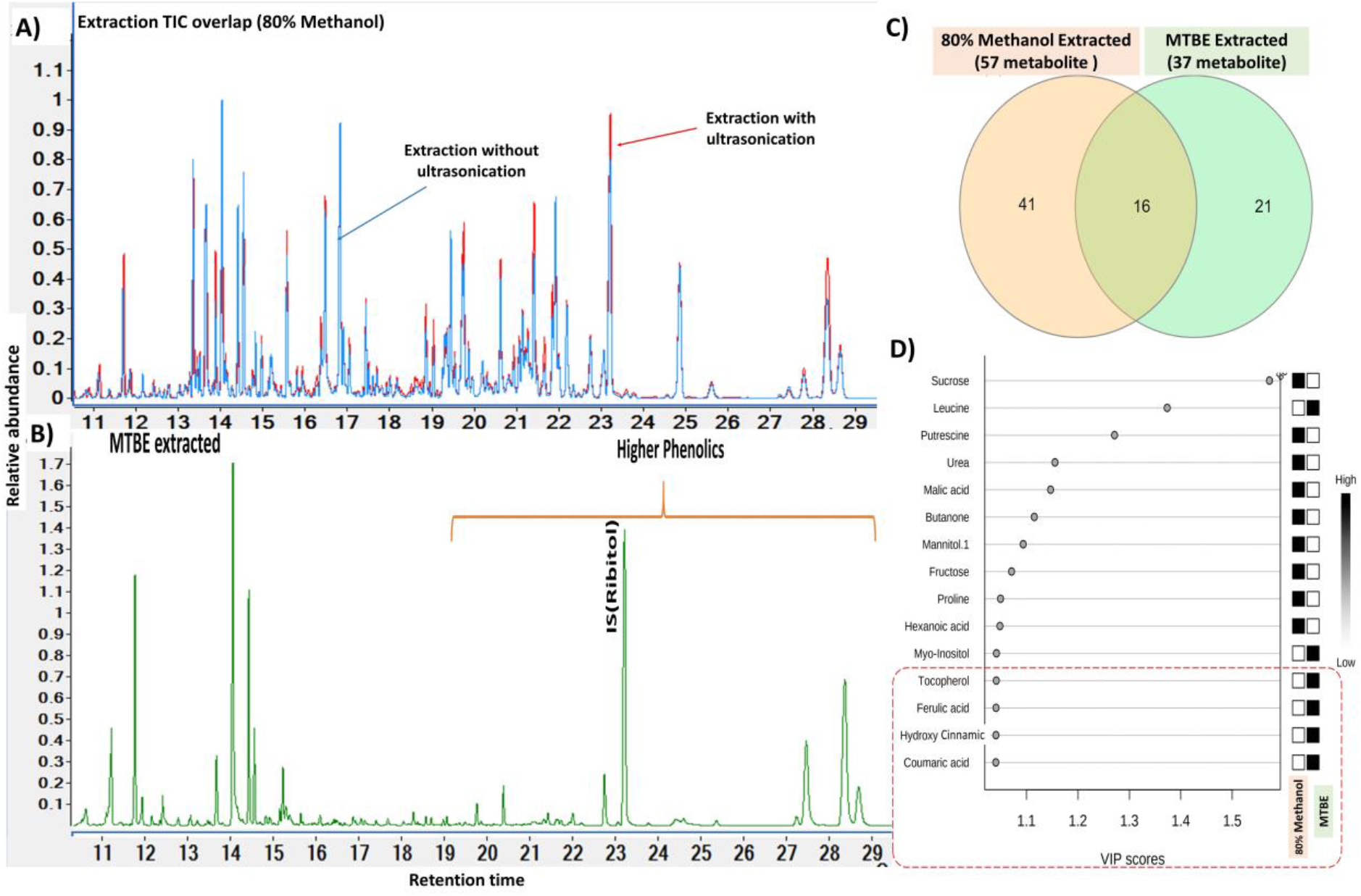
Development and optimization of a robust extraction methodology for phenols. A) Ultrasonication extraction in 80% methanol showed better recovery of phenolics and other metabolites B) MTBE-based phase separation showed better enrichment for phenolics. C) Venn diagram displaying the common and unique metabolites under 80% methanol and MTBE extracts D) Variable Importance in Projection (VIP) score pattern analysis revealed that MTBE recovered phenolics from 80% methanol extracts which primarily showed sugars and organic acids.

### 3.2. Inert N_2_ gas-drying improved stability of the phenols

Phenols are sensitive to light, temperature and oxygen and they may degrade (Ali et al., 2018). We compared drying of extracts under vacuum (speed vac evaporator) and N_2_ purging to find out better techniques for phenolics recovery. Dry N_2_ purging techniques showed higher recovery potential of phenols probably due to the prevention of oxidation loss **(Fig. 4A)**. Both BSTFA and MSTFA derivatisation have shown promising analysis of phenols in GC-MS **(Fig. 4B)**. We further used MSTFA reagent for the derivatisation of plant phenolics.

**Fig. 4.**
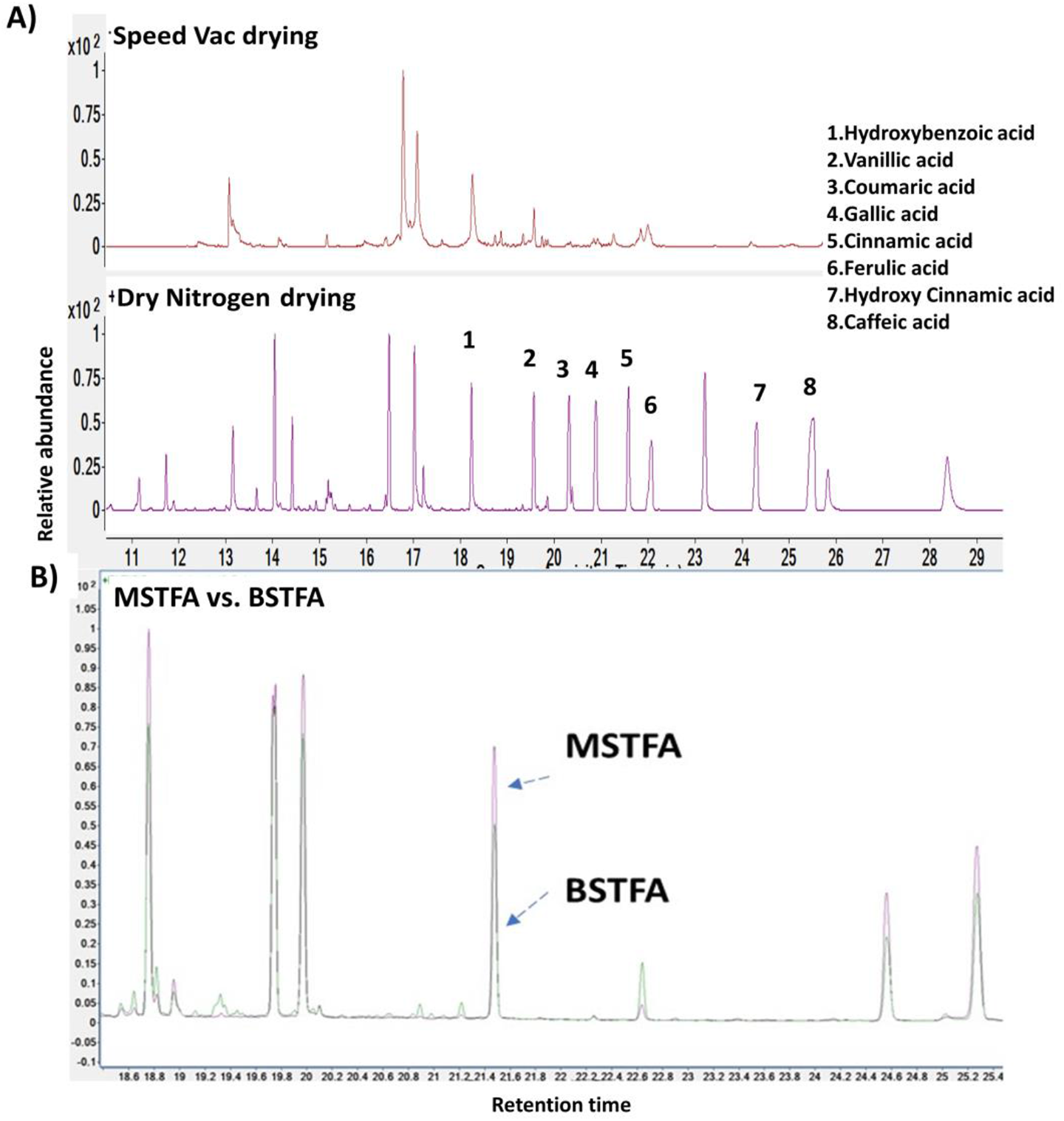
Optimization of comparative drying and derivatisation methodology for phenolics detection. A) Dry N_2_ purging technique showed higher recoveries of 1. Hydroxybenzoic acid, 2. Vanillic acid, 3. Coumaric acid, 4. Gallic acid 5. Cinnamic acid, 6. Ferulic acid 7. Hydroxycinnamic acid and 8. Caffeic acid. B) GC-MS chromatogram showed that MSTFA derivatisation has relatively higher peak abundances of phenolics than BSTFA. The violet spectrum lines represent MSTFA derivatized and the green spectrum lines represents BSTFA derivatized phenolics. The X-axis showed retention times and the Y-axis showed relative peak abundances.

### 3.3. Efficient detection and quantification of Phenolics were achieved using targeted Extracted Ion Chromatograms (EIC) from the GC-MS spectra

In GC-MS analysis, the phenolics were identified on basis of extracted ion fragments, retention time, matching with NIST library and commercial standards. we observed that Extracted Ion Chromatogram (EIC) of targeted m/z of phenolics can reliably be used for quantitative analysis with a better signal vs noise ratio. **(Fig. S2, Fig. S3)**. Phenol standards showed linear response with a promising limit of detection (LOD) (ng/ml) and limit of quantification (LOQ) (ng/ml) as presented in brackets-Cinnamic acid (27.74, 84.06), Coumaric acid (19.03, 57.67), Caffeic acid (20.84, 63.12), Ferulic acid (33.38, 50.57), Hydroxybenzoic acid (27.81, 84.27), Vanillic acid (37.31,113.05), Vanillin (43.55, 131.96), Gallic acid (22.16, 67.14) (**Table 1)**.

**Table 1.**
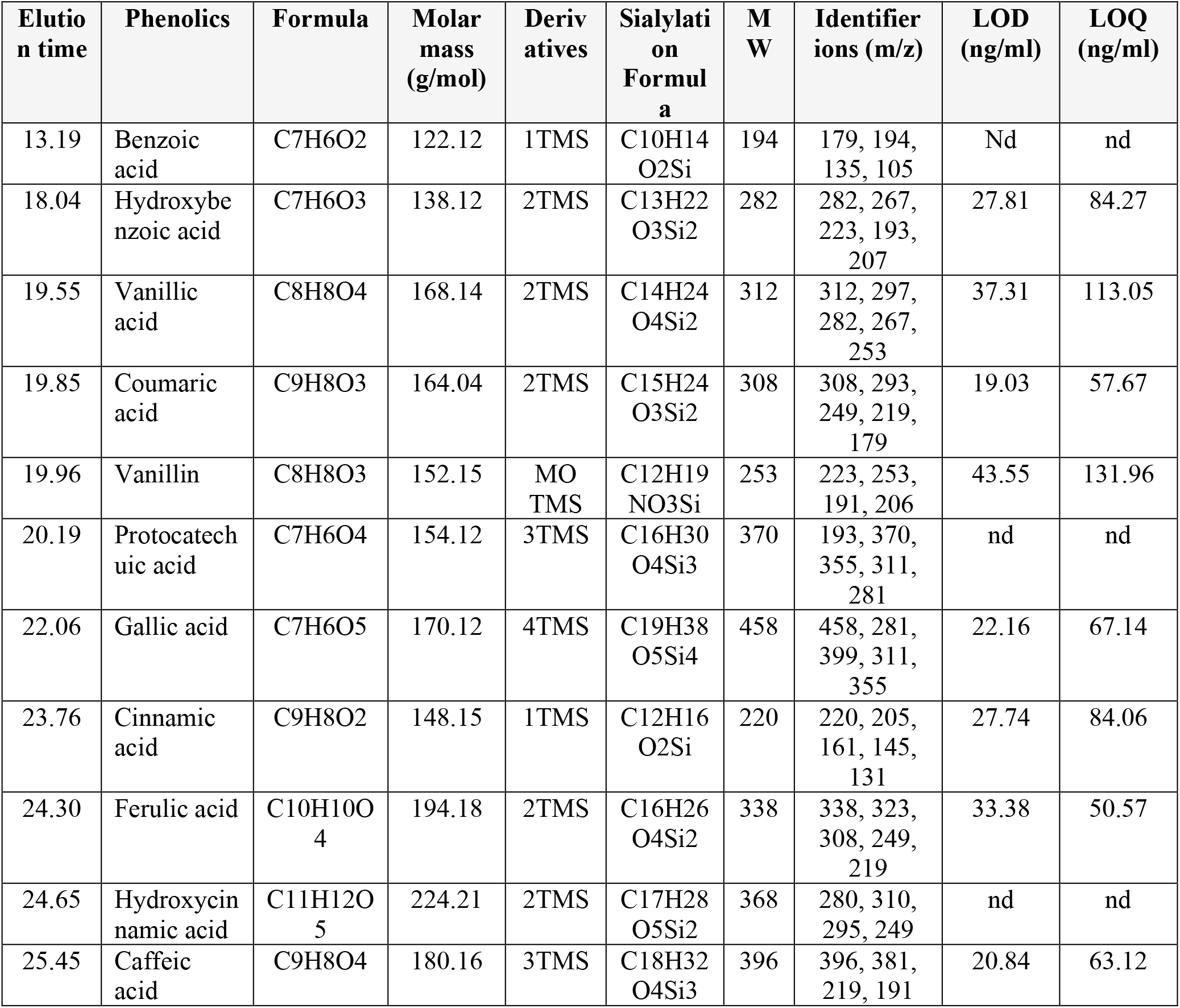
Phenolics detected in *Arabidopsis thaliana* rosette by GC-MS of MTBE extracts. The formula, molar mass, fragments obtained by MSTFA derivatives, identifier ions and their respective LOD and LOQ are listed. A representative GC-MS spectrum of phenolics is provided in supplementary Fig. S2. TMS represent trimethylsilyl, MO represents methyl oxime and “nd” represents Not determined.

| Elution time | Phenolics | Formula | Molar mass (g/mol) | Derivatives | Silylation Formula | MW | Identifier ions (m/z) | LOD (ng/ml) | LOQ (ng/ml) |
| --- | --- | --- | --- | --- | --- | --- | --- | --- | --- |
| 13.19 | Benzoic acid | C<sub>7</sub>H<sub>6</sub>O<sub>2</sub> | 122.12 | 1TMS | C<sub>10</sub>H<sub>14</sub>O<sub>2</sub>Si | 194 | 179, 194, 135, 105 | Nd | nd |
| 18.04 | Hydroxybenzoic acid | C<sub>7</sub>H<sub>6</sub>O<sub>3</sub> | 138.12 | 2TMS | C<sub>13</sub>H<sub>22</sub>O<sub>3</sub>Si<sub>2</sub> | 282 | 282, 267, 223, 193, 207 | 27.81 | 84.27 |
| 19.55 | Vanillic acid | C<sub>8</sub>H<sub>8</sub>O<sub>4</sub> | 168.14 | 2TMS | C<sub>14</sub>H<sub>24</sub>O<sub>4</sub>Si<sub>2</sub> | 312 | 312, 297, 282, 267, 253 | 37.31 | 113.05 |
| 19.85 | Coumaric acid | C<sub>9</sub>H<sub>8</sub>O<sub>3</sub> | 164.04 | 2TMS | C<sub>15</sub>H<sub>24</sub>O<sub>3</sub>Si<sub>2</sub> | 308 | 308, 293, 249, 219, 179 | 19.03 | 57.67 |
| 19.96 | Vanillin | C<sub>8</sub>H<sub>8</sub>O<sub>3</sub> | 152.15 | MO TMS | C<sub>12</sub>H<sub>19</sub>NO<sub>3</sub>Si | 253 | 223, 253, 191, 206 | 43.55 | 131.96 |
| 20.19 | Protocatechuic acid | C<sub>7</sub>H<sub>6</sub>O<sub>4</sub> | 154.12 | 3TMS | C<sub>16</sub>H<sub>30</sub>O<sub>4</sub>Si<sub>3</sub> | 370 | 193, 370, 355, 311, 281 | nd | nd |
| 22.06 | Gallic acid | C<sub>7</sub>H<sub>6</sub>O<sub>5</sub> | 170.12 | 4TMS | C<sub>19</sub>H<sub>38</sub>O<sub>5</sub>Si<sub>4</sub> | 458 | 458, 281, 399, 311, 355 | 22.16 | 67.14 |
| 23.76 | Cinnamic acid | C<sub>9</sub>H<sub>8</sub>O<sub>2</sub> | 148.15 | 1TMS | C<sub>12</sub>H<sub>16</sub>O<sub>2</sub>Si | 220 | 220, 205, 161, 145, 131 | 27.74 | 84.06 |
| 24.30 | Ferulic acid | C<sub>10</sub>H<sub>10</sub>O<sub>4</sub> | 194.18 | 2TMS | C<sub>16</sub>H<sub>26</sub>O<sub>4</sub>Si<sub>2</sub> | 338 | 338, 323, 308, 249, 219 | 33.38 | 50.57 |
| 24.65 | Hydroxycinnamic acid | C<sub>11</sub>H<sub>12</sub>O<sub>5</sub> | 224.21 | 2TMS | C<sub>17</sub>H<sub>28</sub>O<sub>5</sub>Si<sub>2</sub> | 368 | 280, 310, 295, 249 | nd | nd |
| 25.45 | Caffeic acid | C<sub>9</sub>H<sub>8</sub>O<sub>4</sub> | 180.16 | 3TMS | C<sub>18</sub>H<sub>32</sub>O<sub>4</sub>Si<sub>3</sub> | 396 | 396, 381, 219, 191 | 20.84 | 63.12 |

### 3.4. UV-B exposure modulates the levels of free and bounded phenolics in Arabidopsis

Arabidopsis rosette (14 d old) on UV-B treatment showed distinct phenotypic differences. The plants under UV-B treatment (Col-0 UV-B) showed prominent loss of chlorophyll in the leaves in comparison to the healthy and green in the control conditions (Col-0) **(Fig. 5A)**. The UV-B exposed plants, although looked unhealthy they exhibited some tolerance probably due to photoprotective molecules such as phenolics **(Fig. 5B)**. The GC-MS analysis of MTBE extracts of Col-0 and Col-0 UV-B tissues captured 11 phenolics - cinnamic acid, hydroxybenzoic acid, vanillic acid, coumaric acid, gallic acid, ferulic acid, caffeic acid, benzoic acid, hydroxycinnamic acid, vanillin and protocatechuic acid. The fold change analysis and an unpaired t-test revealed that the level of phenolics was significantly enhanced in the Col-0 UV-B plants. Interestingly, coumaric acid (308 m/z, 13.29 folds) and caffeic acid (396 m/z, 12.10 folds) showed very high levels of accumulation (12-14 folds) under UV-B **(Fig. 5C)**. Other phenolics such as cinnamic acid (220 m/z), hydroxybenzoic acid (282 m/z), vanillic acid (312 m/z, gallic acid (458 m/z), ferulic acid (338 m/z), benzoic acid (194 m/z), hydroxycinnamic acid (368 m/z) and protocatechuic acid (370 m/z) also showed elevated levels by about 1 to 4 folds **(Fig. 5C)**. Notably, vanillin (253 m/z) was detected only in the UV-B exposed tissues. Col-0 UV-B also showed enhanced levels of ester-bound phenolics (coumaric acid, gallic acid, hydroxybenzoic acid and vanillic acid), glycoside bound phenolics (vanillic acid and hydroxybenzoic acid) and wall-bounded phenolics (vanillic acid and hydroxybenzoic acid) **(Fig. 6)**. Overall, quantification of phenolics in Col-0 reveals a significant accumulation of coumaric acid, gallic acid, hydroxybenzoic acid, and vanillic acid when treated with continuous UV-B light. The higher levels of phenolics under UV-B exposure seems to reprogram the secondary metabolic pathways leading to the accumulation of free and bounded phenolics in plants for photoprotection and other physiological roles.

**Fig. 5.**
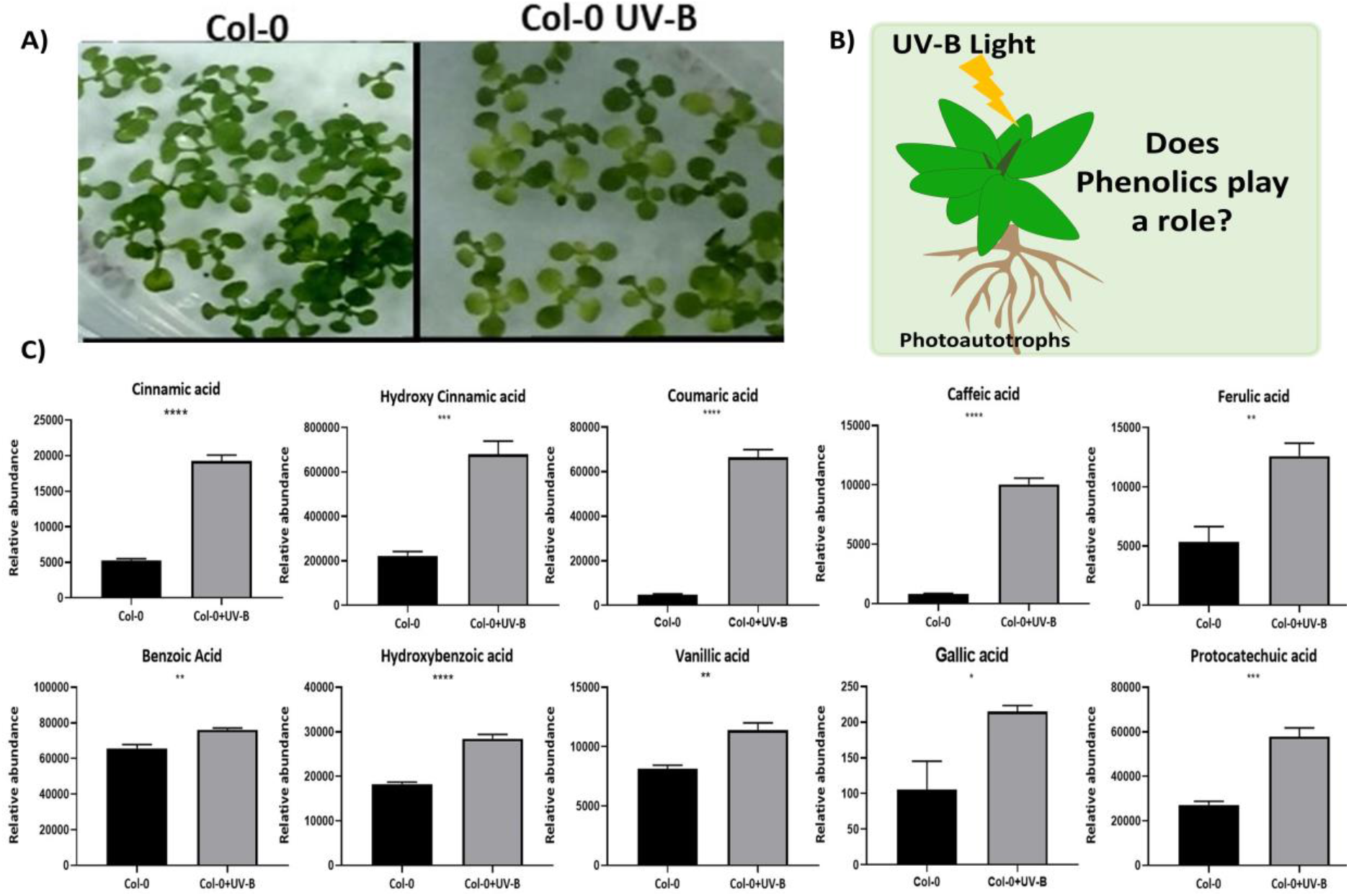
UV-B exposure modulates the level of free phenolics in Arabidopsis. A) The 14 days old Arabidopsis rosette showed distinct phenotypic changes under UV-B as depicted in pictures. B) UV-B showed morpho-physiological adaptation, but we focused to investigate the phenolics of plants because of their photoprotective role. C) GC-MS captured phenolics in Col-0 and Col-0 UV-B. The relative peak abundances of phenolics reveal that the level of phenolic was significantly enhanced under UV-B. Error bars represent the SD of 4 replicates. An unpaired t-test was performed for checking the significance. Asterisks represent statistically significant differences (***P < 0.001, **P < 0.01, *P < 0.05) as determined by t-test.

**Figure 6.**
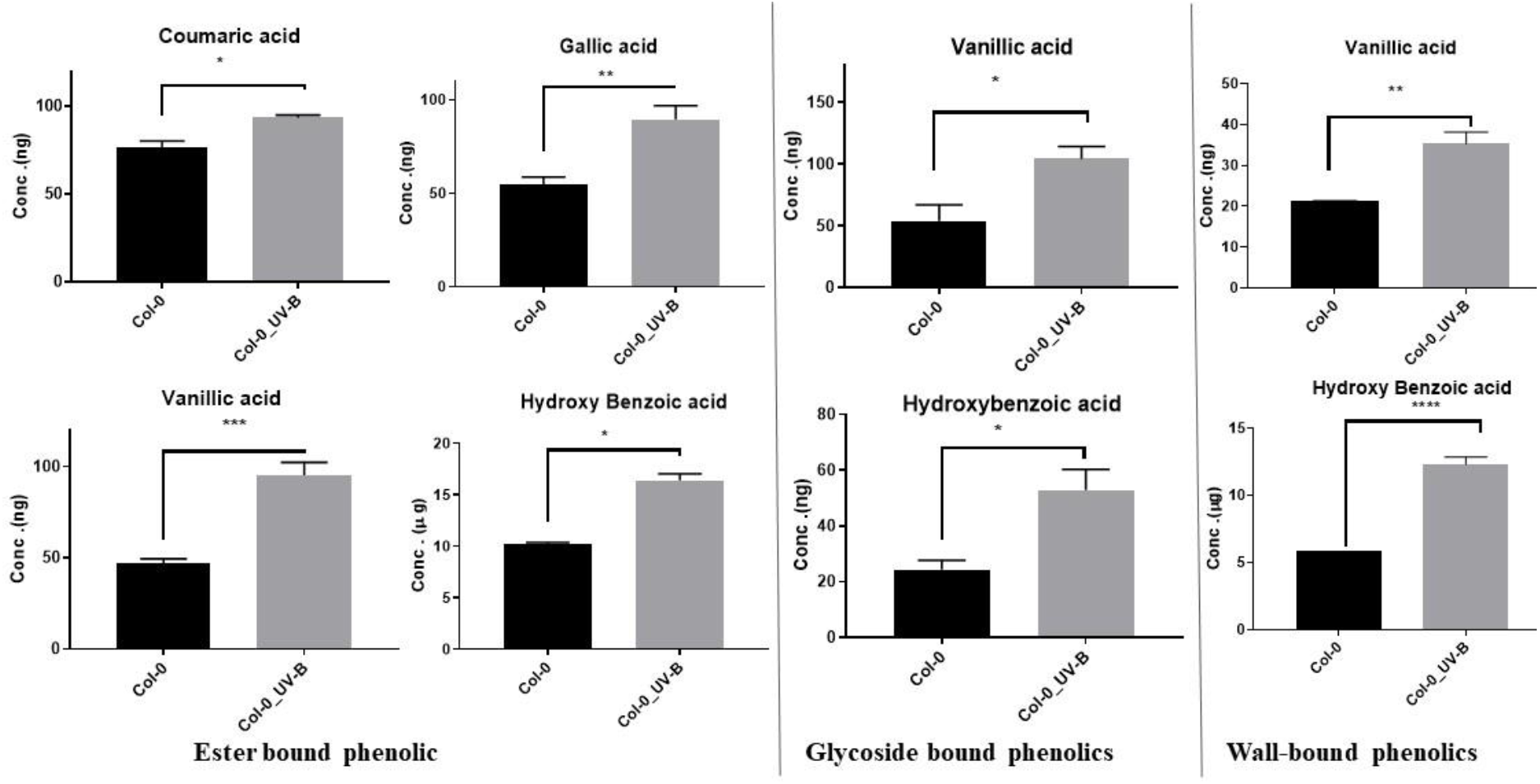
Quantitative levels of Monophenols derived from ester-bound, glycoside bound and wall-bound fractions of Arabidopsis tissues (Col-0 and Col-0_UV-B). UV-B. Monophenols measured are quantified based on per gDW tissues. An unpaired t-test was performed for checking significance, asterisks represent significant differences (***P < 0.001, **P < 0.01, *P < 0.05).

## 4. Discussion

### 4.1. Plant phenolics are ubiquitous and analysing them quantitatively is important

Plant phenolics have relevance to nutrition, health and are known to play significant physiological roles in plants =(Shahidi and Yeo, 2018)(Anantharaju et al., 2016). In plants, the phenolics are biosynthesised via the shikimate pathway and gets accumulated in the cells either as free or as wall-bound in the form of esters or glycosides **(Fig. 1)**. Phenols gain significance in nutrition and health because of their potential antioxidant activities. The levels of phenolics often get modulated in plants depending on their age, environmental factors, biotic or abiotic stress and gene manipulations (Escarpa and Gonzalez, 2001)(Parr and Bolwell, 2000). Qualitative and quantitative analysis of phenolics using silylation and GC-MS has previously been reported (Proestos and Komaitis, 2013). However, these were limited to a number of phenolics, with scope for further improvements in enhancing the range, detection limits and sensitivity. Here we demonstrated a robust method of extraction and GC-MS analysis of Monophenols in plant systems along with a case study which highlighted that UV-B exposure modules the levels of phenols.

### 4.2. Monophenols can effectively be enriched using MTBE solvent, ultrasonication and N_2_ drying

The efficiency of phenols extraction from plant tissues is influenced by a no of factors such as the solvent used and treatment parameters. If not properly optimised, then extraction could compromise the quantitative levels as well as identification of phenolics. Generally, hot aqueous-methanolic solvents are routinely used to extract soluble metabolites which might also contain phenolics (Lisec et al., 2015). But owing to the low abundances of phenols in relation to other soluble central metabolites like sugars, very limited analysis is often possible. The phenols can further be enriched from methanolic extracts by phase separation with other suitable solvents facilitating reduced background noise due to other metabolites. (Gharaati Jahromi, 2019)(Khoddami et al., 2013). Ethyl acetate was previously used for the separation of plant phenolics, but recent research suggested that MTBE can be an alternative suitable solvent. MTBE has high solubility towards analytes and is reported to show homogeneous, clear phase separation and higher extraction recovery during liquid-liquid extraction (Salem et al., 2016)(Juhler and Felding, 2003). Our study established MTBE to be an effective solvent for phase separation that enriched the phenols substantially.

In terms of extraction of phenolics, various methodologies such as solvent extraction, soxhlet extraction, microwave-assisted extraction and ultrasound-assisted extraction showed promise (Dai and Mumper, 2010)(Ehresman, 2003). We found that ultrasonication is low cost, timesaving and is better at the extraction of phenolics from plant tissues. We also observed N_2_ purging can be an adequate drying technology for phenolics, as it prevents oxidative loss of phenols (Lang et al., 2019). In summary, we tested ultrasonication of hot methanolic extracts followed by MTBE phase separation and dried under inert N_2_ gas. The method furnished excellent recovery and sensitivity of monophenolics in GC-MS analysis with LOD of 19 - 44 ng/ml and LOQ of 50 - 132 ng/ml **(Fig. 3, Fig. 4)**.

### 4.3. UV-B stress promotes the accumulation of free and bounded phenolics

Quantification of phenolics as well enhancing their levels in plants that are widely consumed is of immense interest. Here, we applied the robust method of phenol extraction from Arabidopsis tissues under the normal light regime (Col0) and UV-B exposure (Col0-UV-B)and GC-MS analysis. UV-B exposure is known to influence development in plants by damaging photosynthetic machinery and modulates central and secondary metabolism including phenolics (Yadav et al., 2020). The GC-MS analysis of free phenolics reveals that the level of phenolic was significantly enhanced due to UV-B exposure. Also using the optimised protocols, the quantification of ester-bound, glycoside bound and wall-bounded phenolics revealed enhanced accumulation of coumaric acid, gallic acid, hydroxybenzoic acid, and vanillic acid in UV-B **(Fig. 5, Fig. 6)**. UV-B stress promotes the accumulation of free and bounded phenolic compounds in plants, which might help in better survival by conferring photoprotection and other physiological roles. The study highlighted that controlled UV-B exposure might be adopted as a future strategy to enhance phenols in edible plants as a desirable trait.

## 5. Conclusions

Phenolics have vast applications in agriculture, food, nutrition and health. Their efficient extraction, identification and quantification using sensitive analytical platforms are desired. We optimised the analytical and experimental bottlenecks that captured free, ester, glycoside and wall-bound phenolics after acid or alkali treatments of the tissue extracts and subsequent GC-MS analysis. Higher recovery of phenolics was achieved through ultrasonication assisted extraction along with Methyl tert-butyl ether (MTBE) enrichment, nitrogen gas drying and their derivatisation using MSTFA for GC-MS data analysis. The optimised protocol was tested on Arabidopsis rosette exposed to UV-B radiation (280-315 nm) which triggered enhanced levels of 11 phenolics and might be attributed to photoprotection and other physiological roles. Also, the protocol comprehensively can be adopted to capture the variations in the levels of ester, glycoside and wall-bounded phenolics in plant systems under perturbations. The protocol exhibited high reproducibility and sensitivity. The robust method of extraction and GC-MS analysis can widely be adopted for studying phenolics in plant systems.

## Acknowledgements

SKM acknowledges Science and Engineering Research Board (SERB) Early career research funding (File No: ECR/2016/001176). ML acknowledge the Ministry of Education and IIT Mandi for PhD fellowship.

## Supplementary

**Table S1.** GC-MS data acquisition parameters used for profiling of the phenols.

| GC-MS CONTROL PARAMETERS |  |
| --- | --- |
| Injection Source | GC ALS |
| Oven (Run Time) | 60 min |
| Equilibration Time | 0.25 min |
| Max Temperature | 450 °C |
| Temperature (Initial) | 50 °C |
| Hold Time | 5 min |
| Post Run | 70 °C |
| Program |  |
| Oven Program #1 Rate | 10 °C/min |
| Oven Program#1 Value | 200 °C |
| Oven Program#1 Hold Time | 10 min |
| Oven Program#2 Rate | 5 °C/min |
| Oven Program#2 Value | 300 °C |
| Oven Program#2 Hold Time | 10 min |
| Syringe Size | 10 µL |
| Injection Volume | 1 µL |
| Mode | Splitless |
| Liner | Agilent 5190-2295 |
|  | (Split, taper, wool, low pressure drops) |
| Column Information | Agilent 19091S-433, |
|  | HP-5MS 5% Phenyl Methyl Silox |
| Column Temperature Range | 0 °C-325 °C (325 °C) |
| Column Dimensions | 30 m x 250 µm x 0.25 µm |
| Carrier gas flow (Initial) | 0.6 mL/min |
| Gas Flow ,Post Run | 1 mL/min |
| Start Time (M/Z) | 30 |
| Low Mass (M/Z) | 700 |
| MS Source | 230 °C maximum 250 °C |
| MS Quad | 150 °C maximum 200 °C |

**Fig S1.**
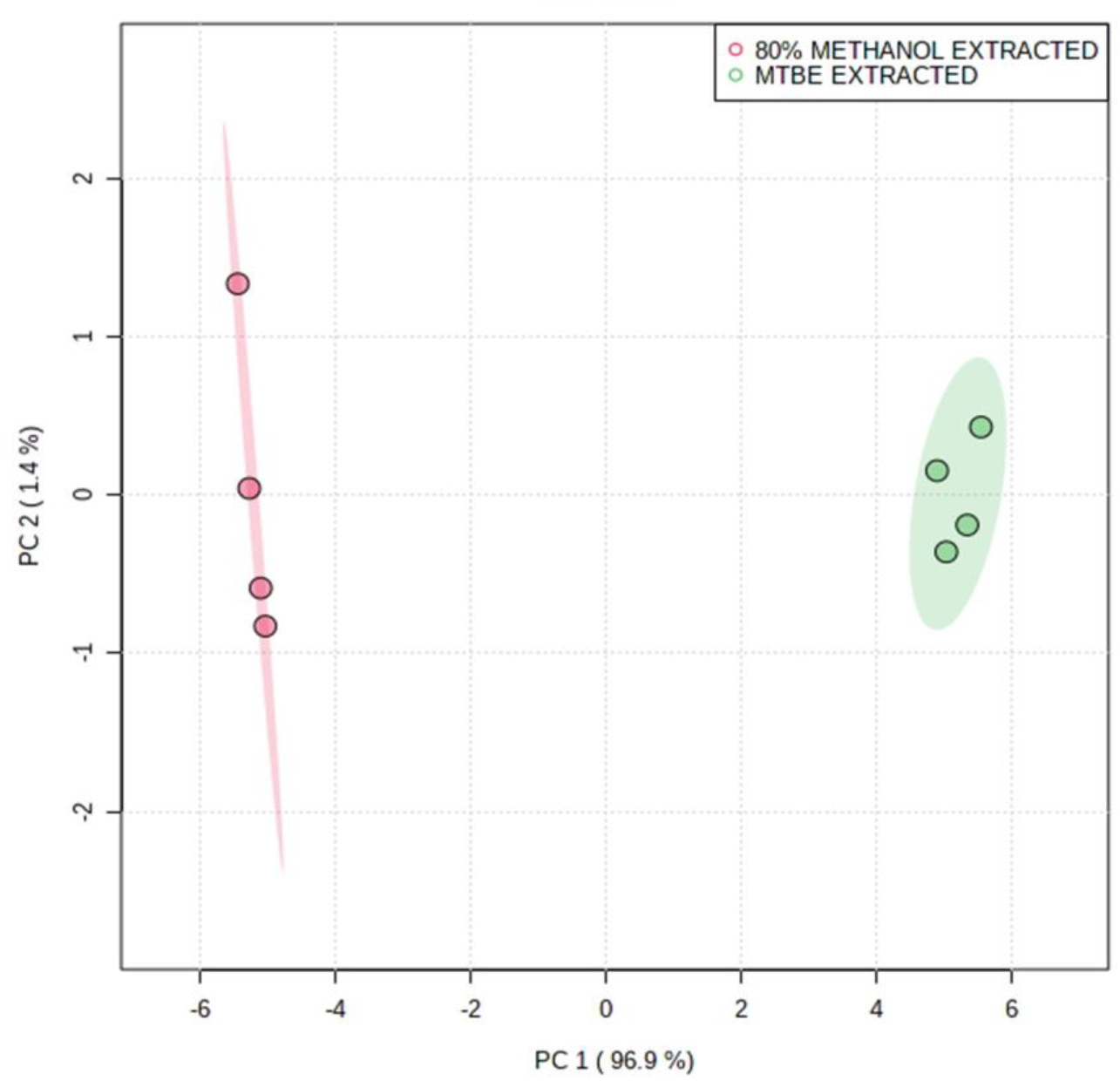
MTBE extraction enriched monophenols predominantly from the methanolic extracts of Arabidopsis tissues. The distinct variations in metabolites are revealed by PCA analysis.

**Fig S2.**
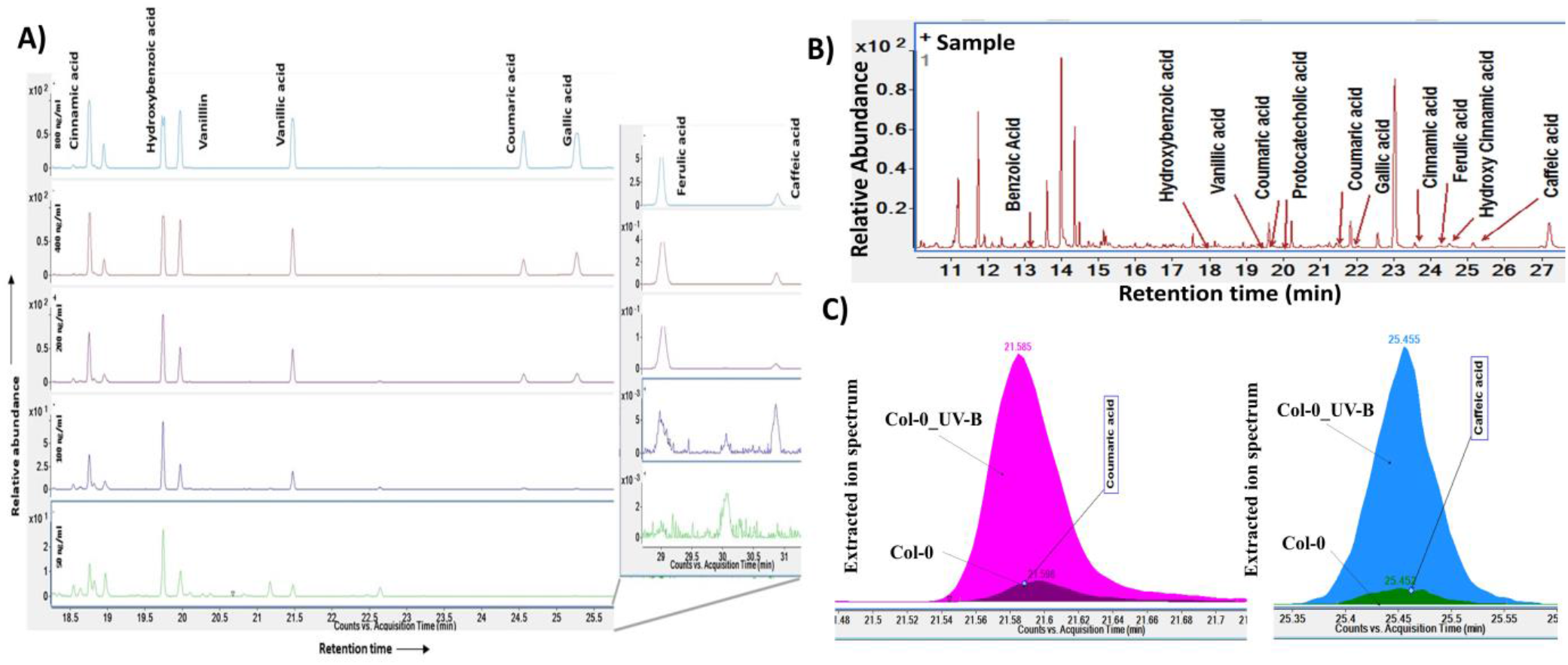
GC-MS analysis of silylated Monophenols A) Monophenol standard mix. B) Representative phenolics detected in Arabidopsis rosette. C) Extracted ion spectrum from Arabidopsis (Col-0 and Col-0_UV-B) reveals UV-B enhanced monophenols

**Fig S3.**
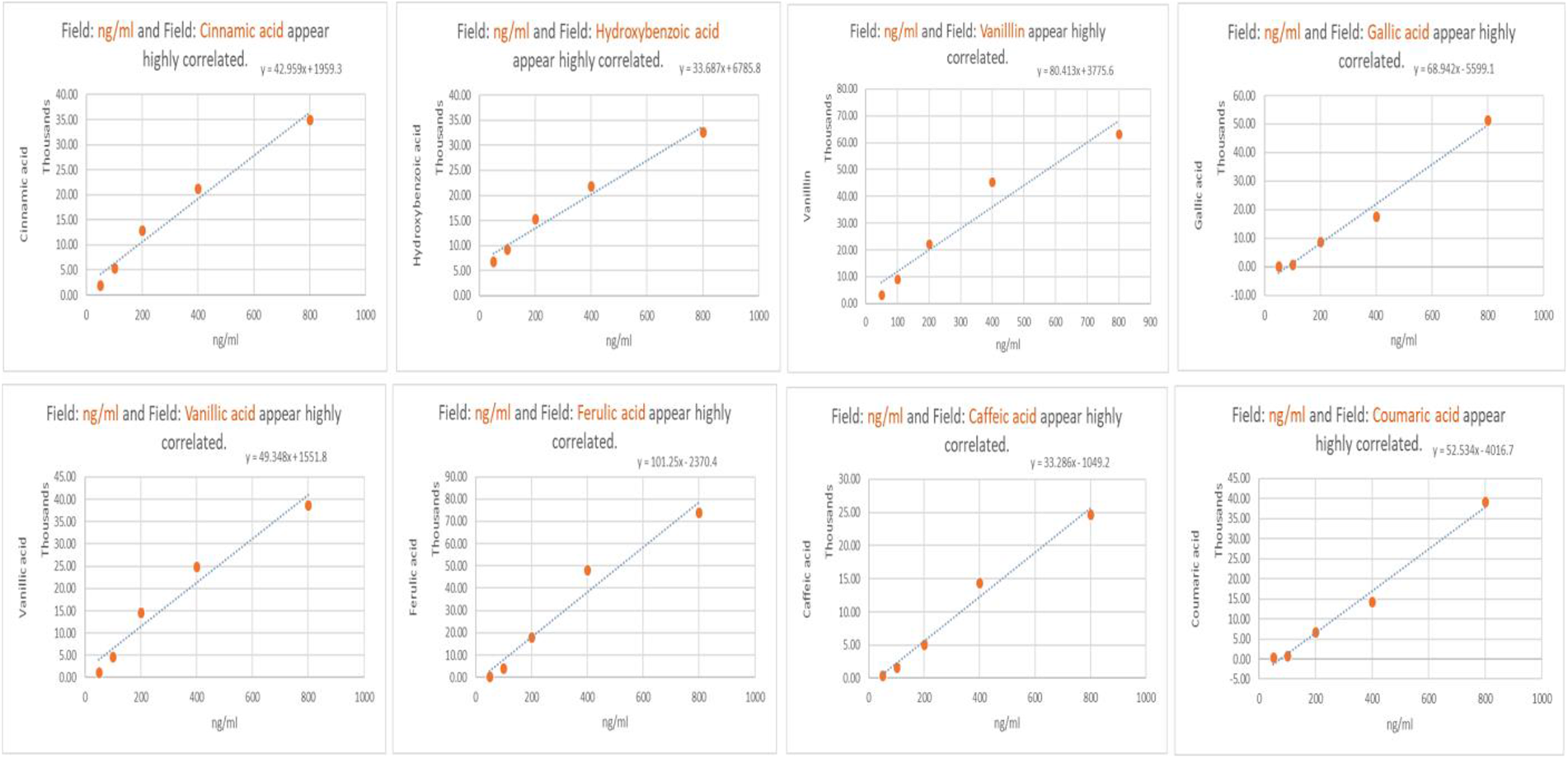
The standard curves of Monophenols based on the abundance of extracted ion chromatograms of GC-MS. These were used to calculate limit of detection (LOD) (ng/ml) and limit of quantification (LOQ) (ng/ml).

**Fig S4.**
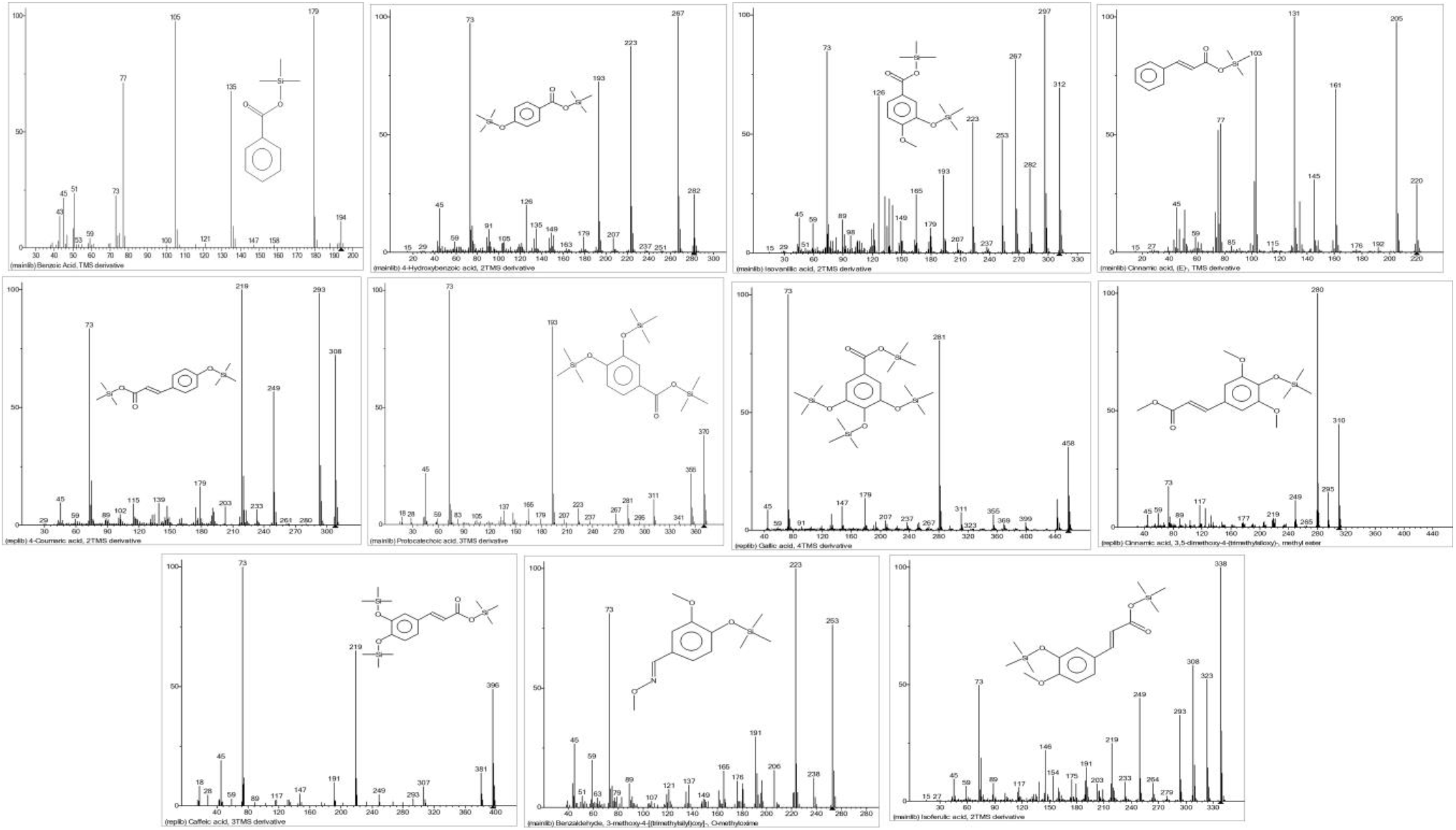
Mass spectra of MSTFA derivatised Monophenols exhibiting the mass ion fragmentation (m/z)

